# Remdesivir inhibits renal fibrosis in obstructed kidneys

**DOI:** 10.1101/2020.04.01.019943

**Authors:** Ming Wu, Lin Xu, Bo Tan, Di Huang, Meijie Yuan, Chaoyang Ye

## Abstract

**Aim:** Kidney impairment is observed in patients with COVID-19. We aimed to demonstrate the effect of anti-COVID-19 agent remdesivir on renal fibrosis.

**Methods:** Remdesivir and its active nucleoside metabolite GS-441524 were used to treat TGF-β stimulated renal fibroblasts (NRK-49F) and human renal epithelial cells (HK2). Cell viability was determined by CCK8 assay, and fibrotic markers were measured by Western blotting. Vehicle or remdesivir were given by intraperitoneal injection or renal injection through the left ureter in unilateral ureteral obstruction (UUO) mice. Serum and kidneys were harvested. The concentrations of remdesivir and GS-441524 were measured using LC-MS/MS. Renal and liver function were assessed. Renal fibrosis was evaluated by Masson’s trichrome staining and Western blotting.

**Results:** Remdesivir and GS-441524 inhibited cell proliferation and the expression of fibrotic markers (fibronectin, pSmad3, and aSMA) in NRK-49F and HK2 cells. Intraperitoneal injection or renal injection of remdesivir attenuated renal fibrosis of UUO kidneys. Renal and liver function were not changed in remdesivir treated UUO mice. Remdesivir can not be detected, but two remdesivir metabolites were detected after injection.

**Conclusion:** Remdesivir inhibits renal fibrosis in obstructed kidneys.

## Introduction

A novel coronavirus (2019-nCoV) reported in Wuhan in late December 2019 has rapidly spread to the rest of the world (1, 2). Novel coronavirus pneumonia is becoming a worldwide public health event due to the rapid increase in new cases and the high severity and mortality (1-3). A latest research shows that COVID-19 patients in the intensive care unit were older (median age 66 vs 51) and more likely to have comorbidities (72.2% vs 37.3%) than patients not in the intensive care unit, suggesting that elderly people or patients with underlying disease have higher disease severity (3). Chronic kidney disease (CKD) is a common disorder and the prevalence of CKD is around 10% in adults (4, 5). Thus, CKD patients combined with COVID-19 should be drawn enough attention from nephrologists.

It is worth noting that the 2019-nCoV can not only cause pneumonia, but also damage other organs, such as the heart, liver and kidneys (6). Angiotensin-converting enzyme 2 (ACE2) mediates the entry of 2019-nCoV coronavirus into human cells (7). It has been found that ACE2 is highly expressed in renal tubular cells, implying that 2019-nCoV may directly bind to ACE2-positive cells in the kidney and thus induce kidney injuries (7). Indeed, a clinical study reported that 27.06% of patients with COVID-19 exhibited acute renal failure (ARF), while elderly patients (≥60 years) were more likely to develop ARF (65.22% vs 24.19%) (8). A further immunohistochemistry analysis revealed that the antigen for 2019-nCoV accumulates in renal tubules (8). Another clinical study with 59 COVID-19 patients showed that proteinuria occurred in 63% of patients (9). 19% and 27% COVID-19 patients have elevated plasma creatinine and urea nitrogen levels, respectively (9). A consecutive cohort study with 710 COVID-19 patients further shows that the prevalence of renal impairment is high, which is associated with in-hospital death (10). Therefore, enough attentions should be paid to kidney protection of COVID-19 patients, and the effect of anti-COVID-19 agents on the kidney should also be concerned.

Renal interstitial fibrosis is a common pathway and main pathological basis for the progression of various chronic kidney diseases to the end-stage renal disease (ESRD) (11, 12). It is characterized by excessive deposition of extracellular matrix in the kidney leading to completely loss of renal function (11, 12). Loss of renal tubule drives the development of renal interstitial fibrosis by producing a large number of profibrotic factors such as TGF-β (12, 13). It has been shown by several animal models that the TGF-β / Smad3 signaling pathway play a key role in renal fibrosis (14, 15).

Remdesivir (GS-5734) is a nucleoside analogue designed for the treatment of severe acute respiratory syndrome coronavirus (SARS), the Middle East respiratory syndrome (MERS) and Ebola virus (16, 17). It can be anabolized to the active triphosphate metabolite and then incorporated into the newly synthesized RNA strand of the virus as a substrate for viral RNA-dependent RNA synthetase (RdRp), thereby prematurely terminating viral RNA transcription (16, 17). *In vitro* study shows that remdesivir can effectively inhibit the infection of 2019-nCoV (18). A single case study shows that treatment with remdesivir improved the clinical condition of the first severely infected COVID-19 patient in the United States in 24 hours (19). Currently, several clinical trials using remdesivir as a treatment for infected COVID-19 patients are undergoing around the world. The first phase III clinical trial which is conducted in China is expected to be completed in the end of April 2020 (ClinicalTrials.gov Identifier: NCT04257656).

The aim of the current study is to determine the effect remdesivir on renal fibrosis.

## Results

### Remdesivir inhibited renal fibrosis *in vitro*

GS-441524, the active metabolite of remdesivir, was used to treat TGF-β stimulated rat renal interstitial fibroblasts (NRK-49F) and huamn renal epithelial cells (HK2) (20). 24 hours stimulation with 2.5 ng/ml TGF-β increased the cell proliferation of NRK-49F cells as shown by CCK8 assay, and treatment with the highest concentration (100 μM) of GS-441524 significantly inhibited its cell proliferation (Figure 1A). There was no dead cells and cell morphology was normal at the highest concentration (100 μM) of GS-441524 when observed by microscopy.

**Figure 1.**
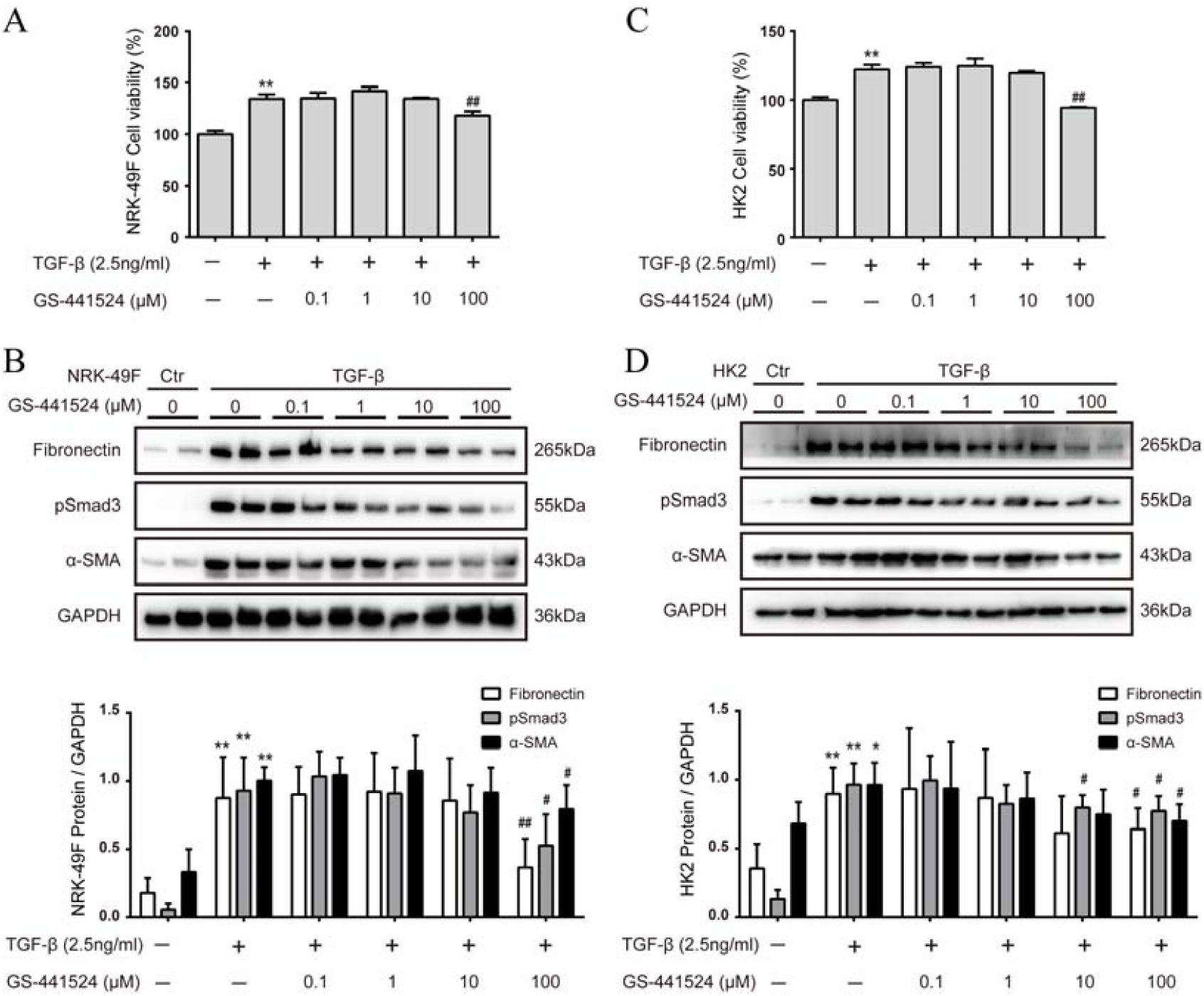
GS-441524, the intermediate metabolite of remdesivir, inhibited renal fibrosis *in vitro*. (A) Following TGF-β stimulation, renal fibroblasts NRK-49F cells were treated with various concentration (0, 0.1, 1, 10, 100 μM) of GS-441524 for 24 hours. CCK8 assay was performed to determine the viability of NRK-49F cells. (B and C) The expression of fibronectin (FN), phosphorylated Smad3 (pSmad3), and alpha-smooth muscle actin (α-SMA) of NRK-49F cells were analyzed by Western blotting and quantified; (D) Upon TGF-β stimulation, human renal epithelial HK2 cells were treated with various concentration (0, 0.1, 1, 10, 100 μM) of GS-441524 for 48 hours. CCK8 assay was performed to determine the viability of HK2 cells. (E and F) The expression of FN, pSmad3, and α-SMA of HK2 cells were analyzed by Western blotting and quantified. Data represent mean ± SD. \**P*< 0.05 versus Vehicle-DMSO; \*\**P*< 0.01 versus Vehicle-DMSO; #*P*< 0.05 versus TGF-β-DMSO; ##*P*< 0.01 versus TGF-β-DMSO. One representative result of at least three independent experiments is shown.

The protein expression of fibronectin (FN), phosphorylation of Smad3 (pSmad3), and alpha smooth muscle actin (α-SMA) were assessed by Western blotting as markers for fibrosis. TGF-β stimulation increased the expression of FN, pSmad3 and α-SMA in NRK-49F cells, and dose-dependent inhibition of these fibrotic markers was observed starting from 10 μM of GS-441524 (Figure1B).

The effect of GS-441524 on renal fibrosis was further studied in renal epithelial cells (HK2). 48 hours stimulation with 2.5 ng/ml TGF-β increased the cell proliferation of HK cells as shown by CCK8 assay, and inhibition of cell proliferation was observed at 100 μM of GS-441524 (Figure 1C). Remarkable inhibition of FN, pSmad3, and α-SMA expression by GS-44152 was observed starting from 10 μM of GS-441524 (Figure 1D).

We further tested the direct effect of remdesivir *in vitro*. 24 hours treatment with remdesivir slightly inhibited cell proliferation of TGF-β stimulated NRF-49F cells at the highest concentration (10 μM), although there was no significant difference (Figure 2A). Inhibition of FN, pSmad3, and α-SMA expression by remdesivir was observed starting from 1 μM of GS-441524 (Figure 2B).

**Figure 2.**
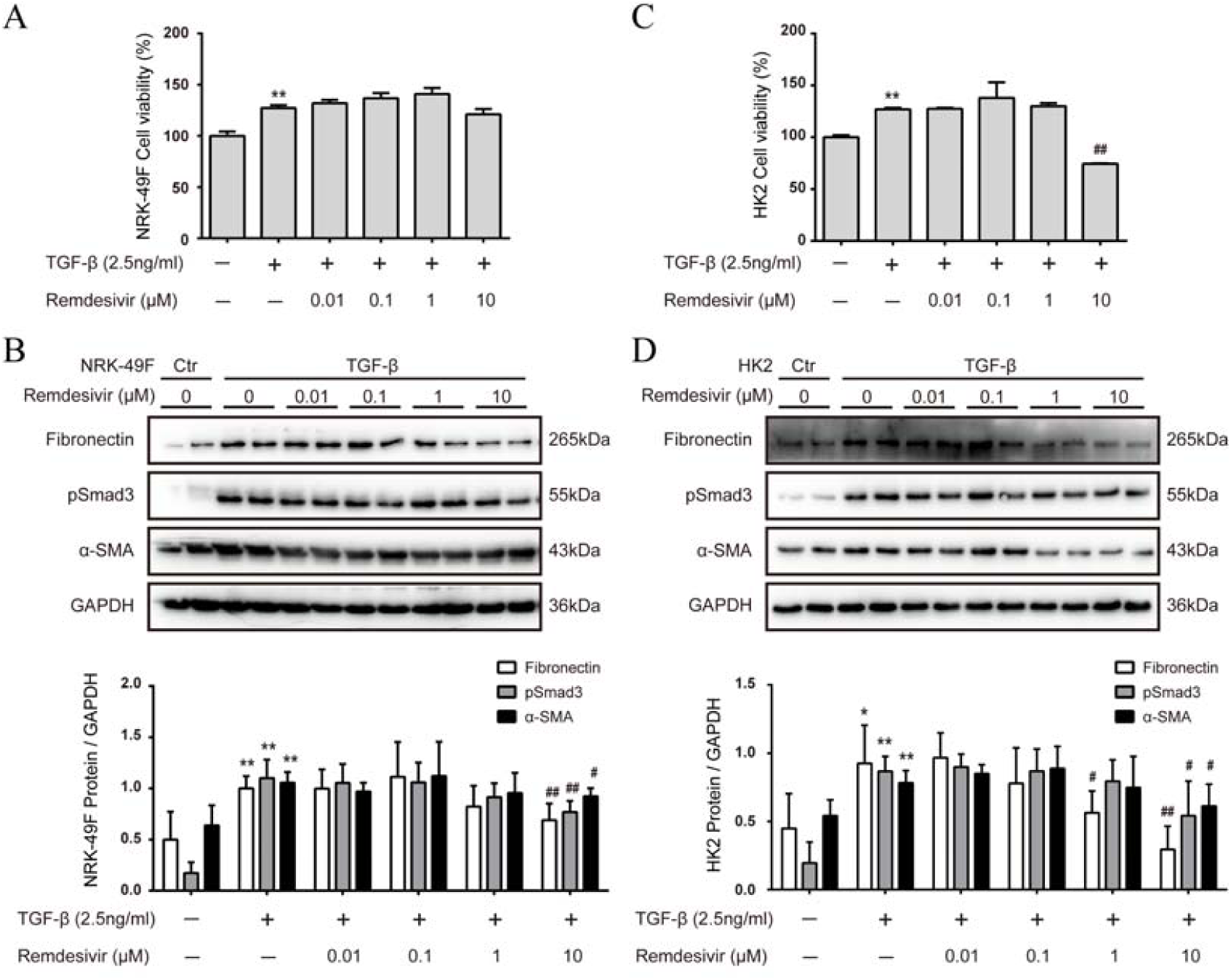
Remdesivir inhibited renal fibrosis *in vitro*. (A) Following TGF-β stimulation, renal fibroblasts NRK-49F cells were treated with various concentration (0, 0.01, 0.1, 1, 10 μM) of remdesivir for 24 hours. CCK8 assay was performed to determine the viability of NRK-49F cells. (B and C) The expression of FN, pSmad3, and α-SMA of NRK-49F cells were analyzed by Western blotting and quantified; (D) Upon TGF-β stimulation, human renal epithelial HK2 cells were treated with various concentration (0, 0.01, 0.1, 1, 10 μM) of remdesivir for 48 hours. CCK8 assay was performed to determine the viability of HK2 cells. (E and F) The expression of FN, pSmad3, and α-SMA of HK2 cells were analyzed by Western blotting and quantified. Data represent mean ± SD. \**P*< 0.05 versus Vehicle-DMSO; \*\**P*< 0.01 versus Vehicle-DMSO; # *P*< 0.05 versus TGF-β-DMSO; ##*P*< 0.01 versus TGF-β-DMSO. One representative result of at least three independent experiments is shown.

Figure 2C shows that 48h treatment with remdesivir significantly inhibited cell proliferation at the highest concentration of remdesivir (10 μM), and normal cell morphology was observed at this concentration. The expression of FN, pSmad3, and α-SMA were significantly down-regulated by remdesivir starting from 1 μM (Figure 2D).

### Intraperitoneal injection of remdesivir inhibited renal fibrosis in UUO mice

Mouse renal fibrosis model was established by UUO operation. One day after sham or UUO operations, mice were treated with vehicle or remdesivir for 10 days. Treatment with remdesivir had no effect on body weight of sham and UUO mice, and all mice were survived during the treatment (data not shown). Mild interstitial fibrosis was observed in vehicle treated UUO mice, which was attenuated by remdesivir (Figure 3A). The protein expression of FN, collagen-I (Col-I), pSmad3, and α-SMA were up-regulated in UUO mouse kidneys as compared with that in sham operated mouse kidneys, and the treatment with remdesivir significantly reduced the expression of these pro-fibrotic proteins in UUO mouse kidneys (Figure 3B). Liver function (ALT and AST) and renal function (Scr and BUN) were determined. Remdesivir has no effect on either liver function or renal function (Figure 3C). Serum and kidney concentration of remdesivir and two metabolites of remdesivir (GS-441524 and Ala-Met) were determined by LC-MS/MS. Remdesivir can not be detected in serum or kidney in both sham and UUO mice (data not shown). However, GS-441524 can only be detected in the serum and kidney of remdesivir treated sham or UUO mice (Figure 3D). Similarly, serum or kidney Ala-Met can only be detected in remdesivir treated sham or UUO mice (Figure 3D).

**Figure 3.**
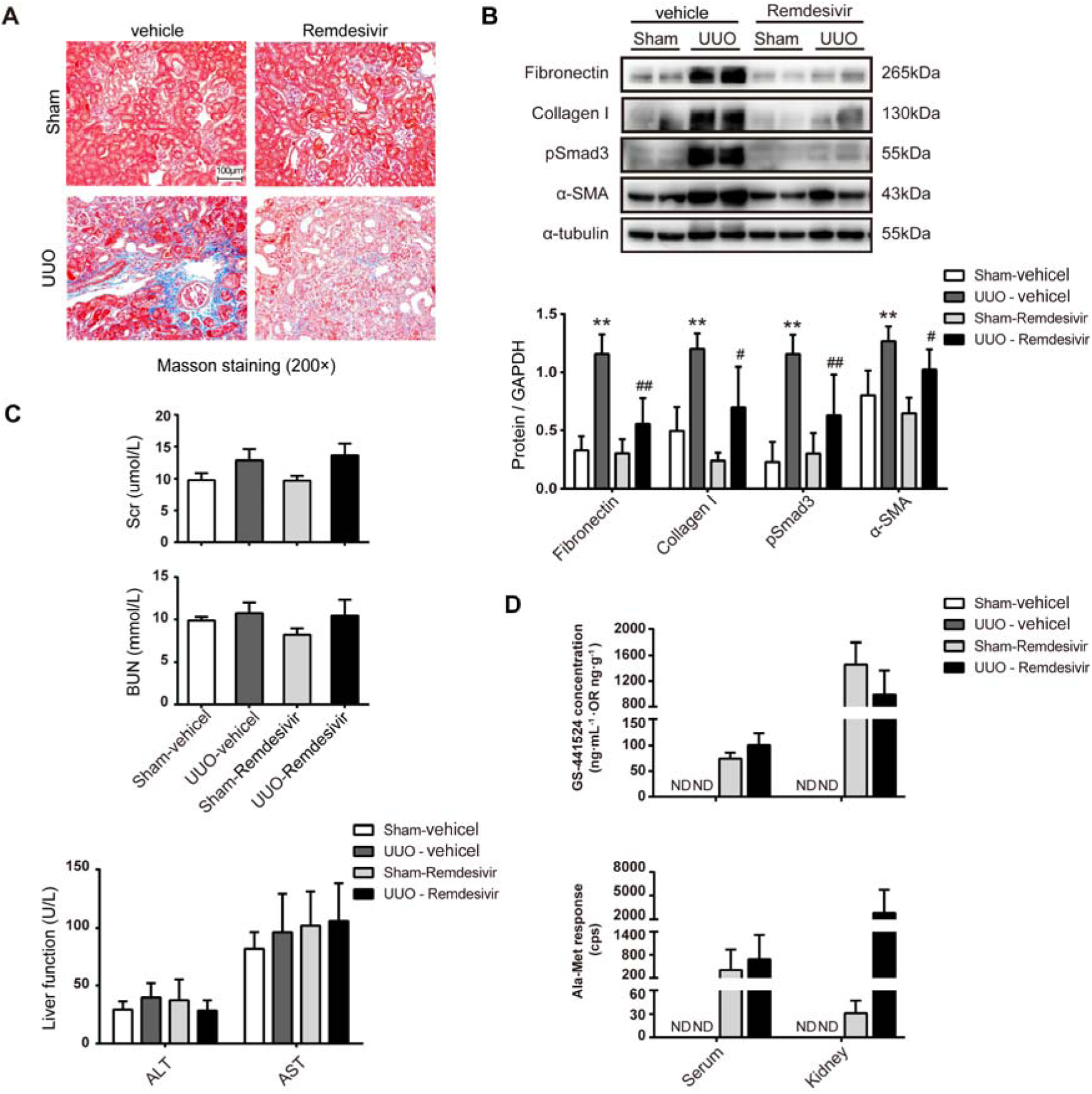
Intraperitoneal (i.p.) administration of remdesivir inhibited fibrosis in UUO mice. After sham or UUO operation, wide type c57 mice were treated with 10 mg/kg/d remdesivir by i.p. injection for 10 days. Serum and kidney tissue were collected 1 hour after remdesivir injection at day 10. (A) Renal fibrosis was assessed by Masson’s trichrome staining. (B) The expression of FN, collagen I (Col-1), pSmad3, and α-SMA by Western blotting. One representative of at least three independent experiments is shown. (C) Liver function (ALT and AST) and renal function (Scr and BUN) were assessed. (D) The concentrations of nucleoside metabolite (GS-441524) and alanine metabolite (Ala-Met), two remdesivir metabolites, in serum and kidneys were determined by LC-MS/MS. Data represent mean ± SD. ND represents not determined. \**P*< 0.05 versus Sham-vehicle; \*\**P*< 0.01 versus Sham-vehicle; #*P*< 0.05 versus UUO-vehicle; ##*P*< 0.01 versus UUO-vehicle.

### Renal injection of remdesivir inhibited renal fibrosis in UUO mice

Remdesivir (1 mg/mL, 50 μl/mouse) or vehicle was infused retrogradely through ureter to the left kidney which was subjected to unilateral utero ligation (UUO) operation. Masson staining shows that interstitial fibrosis was attenuated in UUO kidneys at day7 by local remdesivir injection (Figure 4A). The expression of FN, Col-I, pSmad3, and α-SMA were reduced in remdesivir treated UUO kidneys as compared with that in vehicle treated control kidneys at day7 as shown by Western blotting (Figure 4B). Liver function (ALT and AST) and renal function (Scr and BUN) were not changed by remdesivir in UUO mice (Figure 4C). The remdesivir metabolite GS-441524 can only be detected in serum and kidney at 1 hour after injection only in remdesivir treated mice, which can not be detected at day7 (Figure 4D). The Ala-Met is abundant in remdesivir treated kidneys at 1 hour after injection and it was reduced to background level at day7 (Figure 4D).

**Figure 4.**
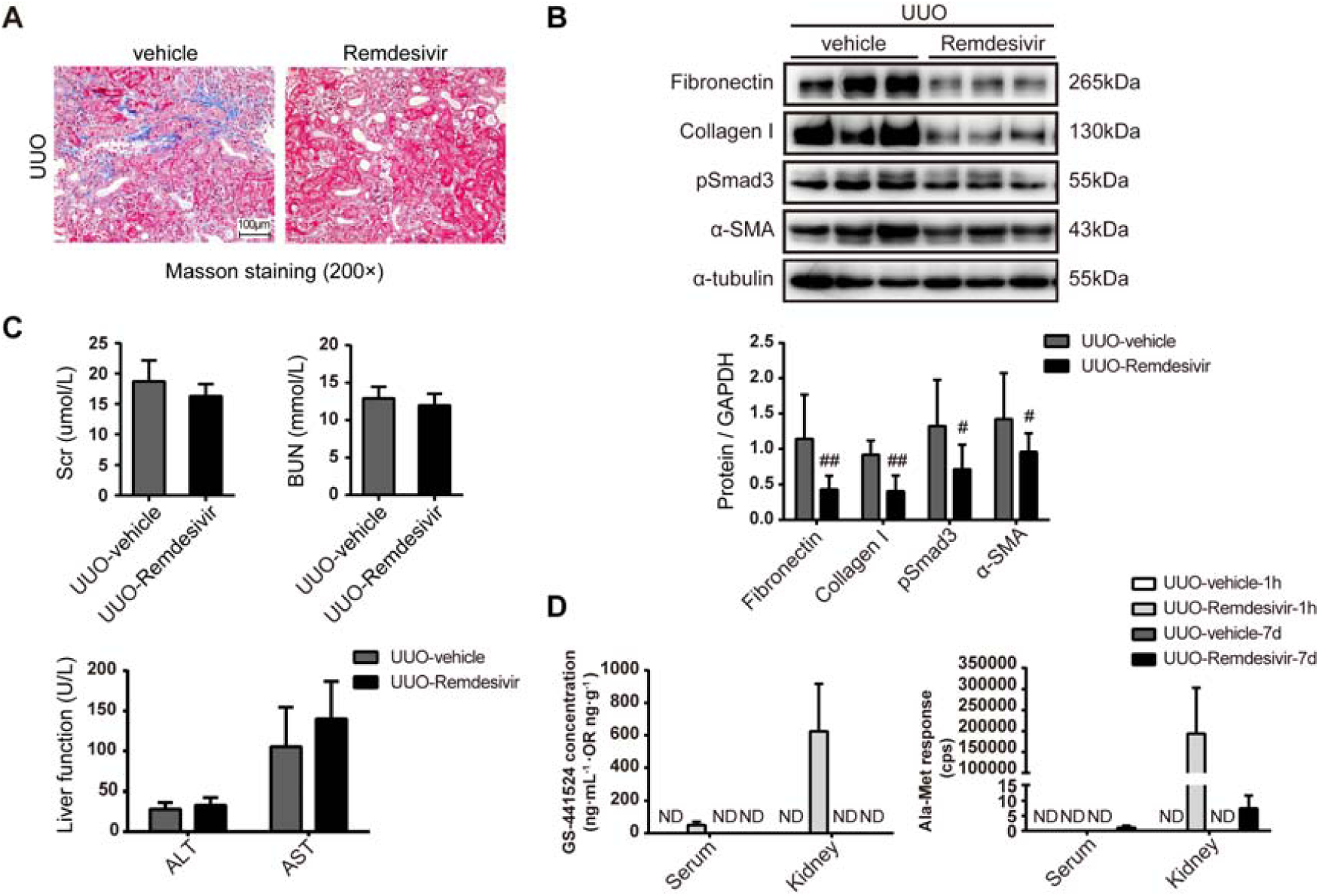
Intrarenal administration of remdesivir inhibited fibrosis in UUO mice. 50μL of vehicle or remdesivir (1 mg/mL) was injected intrarenally to the left kidney, which was subjected to the UUO operation thereafter. Serum and renal tissues were collected at 1 hour or day7 after UUO operation. (A) Renal fibrosis was assessed by Masson’s trichrome staining. (B) The expression of FN, collagen I (Col-1), pSmad3, and α-SMA by Western blotting. One representative of at least three independent experiments is shown. (C) Liver function (ALT and AST) and renal function (Scr and BUN) were assessed. (D) The concentrations of nucleoside metabolite (GS-441524) and alanine metabolite (Ala-Met), two remdesivir metabolites, in serum and kidney were determined by LC-MS/MS. Data represent mean ± SD. ND represents not determined. #*P*< 0.05 versus UUO-vehicle; ##*P*< 0.01 versus UUO-vehicle.

## Discussion

The effect of remdesivir on renal fibrosis is currently unknown. In the present study, we showed that remdesivir inhibits renal fibrosis. First, treatment with remdesivir or its active metabolite GS-441524 inhibited fibrosis in two cellular models for renal fibrosis. Second, systematic administration of remdesivir inhibited renal fibrosis as shown by Masson staining and Western blotting. Third, local infusion of remdesivir into UUO kidneys further revealed the anti-fibrotic effect of remdesivir.

The limitation of this study is that UUO model is not suitable to study the pharmaceutical effect on renal function. UUO model is a classic model to study renal fibrosis, although we did observe a slightly increase in Scr and BUN levels in UUO mice as compared with that in sham mice (21). The renal and liver function were assessed in this study to exclude the potential toxic effect of remdesivir, and we show no effect of remdesivir on renal and liver function in this study.

The concentration of remdesivir was measured to assess the dosing efficacy. However, we are not able to detect remdesivir in serum or kidneys in two different dosing regimens or at different time points. This is probably due to the rapid turnover of remdesivir by esterases which are abundant in blood and tissues (17, 22). Next, we measured the concentration of two remdesivir metabolites, alanine metabolite (Ala-Met) and nucleoside metabolite (GS-441524) (17). GS-441524 and Ala-Met can be detected at 1 hour after IP injection in serum and kidneys of sham and UUO mice. GS-441524 was detected in the serum and kidney at 1 hour after renal injection but not at day 7 post-injection. To prove the specificity of renal infusion of remdesivir, we further measured Ala-Met which can only be detected in kidneys but not in serum at 1 hour after renal injection, and it was not detectable at day 7 after renal injection.

In conclusion, we showed that remdesivir inhibits renal fibrosis in obstructed kidneys.

## Methods

### Animals and UUO operation

Male C57 mice (SPF grade, 20-25g) were purchased and housed in Shanghai Model Organisms Center Inc (SMOC) according to local regulations and guidelines.

After anesthesia with sodium pentobarbital (8mg/kg, i.p.), the left mouse kidney was exposed by an incision. UUO operation was performed through twice ligation of the left ureter with 4-0 nylon sutures. Animal experiments described here in were approved by the animal experimentation ethics committee of Shanghai University of Traditional Chinese Medicine (PZSHUTCM18111601).

For the experiment of 10 days treatment by intraperitoneal (i.p.) injection, mice were randomly divided into four groups: (I) Sham + vehicle (n=5), (II) Sham + remdesivir (n=7), (III) UUO + vehicle (n=8), and (IV) UUO + remdesivir (n=8) group. Mice were treated with vehicle or 10 mg/kg remdesivir daily by i.p. injection. Mice were sacrificed at day 10 at 1 hour after last injection. Serum and kidney tissue were collected.

For the experiment of 1 hour or 7 days treatment by intrarenal injection, mice were randomly divided into two groups: (I) UUO + vehicle (n=11), and (II) UUO + remdesivir (n=11) group. Four mice from each group were sacrificed at 1 hour after renal injection, and the rest of mice were sacrificed at day7. Serum and kidney tissue were collected. Alanine transferase (ALT), aspartate aminotransferase (AST), blood urea nitrogen (BUN) and serum creatinine (Scr) values were assessed in clinical laboratory of Shuguang hospital using a routine method.

### In Vivo Drug Administration

Remdesivir (Product name GS-5734; Cat. No. CSN19703) was purchased from CSNpharm (Chicago, Illinois, USA) and dissolved in DMSO as a 50mg/ml stock, which was further diluted into normal saline by sonication as a working solution. 0.04% typan blue dye (A601140, Sangon, Shanghai, China) was added into vehicle or remdisivir working solution to monitor the injection process. 50 μL of vehicle or remdesivir (1mg/mL) was injected retrogradely once into the left kidney via the ureter. Right after the injection, unilateral ureteral obstruction was performed.

### Cell culture

HK2 renal proximal tubular epithelial cells were obtained from the Cell Bank of Shanghai Institute of Biological Sciences (Chinese Academy of Science). NRK-49F rat kidney interstitial fibroblast cells were purchased from National Infrastructure of Cell Line Resource, Chinese Academy of Medical Sciences. HK2 and NRK-49F cells were cultured in DMEM/F12 medium containing 10%FBS and 0.5% penicillin and streptomycin in an atmosphere of 5% CO2 and 95% air at 37°C. For Western blotting, HK2 and NRK-49F cells were seeded in 6-well plate to 40-50% confluence, which were starved with DMEM/F12 medium 0.5% fetal bovine serum overnight before the experiment. In the next day, fresh medium containing 0.5% fetal bovine serum was changed, and then cells were exposed to 2.5 ng/ml TGF-β (Peprotech, Rocky Hill, NJ, USA) for 24hr or 48h in the presence of various concentration of GS-441524 (Catalog No. T7222, Targetmol, Boston, MA, USA) or remdesivir (Product name GS-5734; Cat. No. CSN19703; CSNpharm, Chicago, IN, USA).

For the CCK8 assay, HK2 (15-25% confluence) and NRK-49F (30-40% confluence) cells were seeded in tetraplicates in 96-well plates, which were starved with DMEM/F12 medium 0.5% fetal bovine serum overnight before the experiment. In the next day, fresh medium containing 0.5% fetal bovine serum was changed, and then cells were exposed to 2.5 ng/ml TGF-β for 24hr-48hr in the presence of various concentration of GS-441524 or remdesivir. After that, 10 μl CCK8 (YEASEN, Shanghai, China) solution was added to each well and cells were incubated for 1 hr. Cell viability was assessed by measure the absorbance at 450 nm.

### Quantitation of remdesivir and its two metabolites

Remdesivir and its two metabolites, alanine metabolite (Ala-Met) and nucleoside metabolite (GS-441524), in serum and kidney were determined using a LC-MS/MS method as described in a previous literature with minor revision (17). In brief, 200 ul of serum or kidney homogenates were mixed with equivalent volume of acetonitrile-methanol mixture (1: 1, v/v). Then, internal standards were added, vortexed, and centrifuged at 15,000 g for 5 minutes. The supernatant was collected and mixed with equivalent volume of deionized water. An aliquot of 10 μl was subsequently injected into a Waters LC-MS/MS system which contains an ACQUITY UPLC and a Xevo TQ-S tandem quadrupole mass spectrometry (Waters, Milford, MA, USA). Standard solutions of remdesivir and GS-441524 were used to plot calibration curves for quantification. Due to the standard is commercially unavailable, alanine metabolite was semi-determined by mass spectrometry response.

### Masson’s trichrome

Mouse kidneys were fixed in 4% paraformaldehyde and further embedded in paraffin. Masson’s trichrome staining was performed using a standard protocol. Briefly, the Four-μm-thick sections of paraffin-embedded kidney tissue was stained with hematoxylin, and then with ponceau red liquid dye acid complex, which was followed by incubation with phosphomolybdic acid solution. Finally, the tissue was stained with aniline blue liquid and acetic acid. Images were obtained with the use of a microscope (Nikon 80i, Tokyo, Japan).

### Western blotting analysis

Renal protein was extracted from the medulla and cortex of mouse kidneys. The protein concentration was measured by the Bradford method, and the supernatant was dissolved in 5x SDS-PAGE loading buffer (P0015L, Beyotime Biotech, Nantong, China). Samples were subjected to SDS-PAGE gels. After electrophoresis, proteins were electro-transferred to a polyvinylidene difluoride membrane (Merck Millipore, Darmstadt, Germany), which was incubated in the blocking buffer (5% non-fat milk, 20mM Tris-HCl, 150mMNaCl, PH=8.0, 0.01%Tween 20) for 1 hour at room temperature and was followed by incubation with anti-fibronectin (1:1000, ab23750, Abcam), anti-pSmad3 (1:1000, ET1609-41, HUABIO), anti-Collagen I (1:500, AF7001, sc-293182, Santa Cruz), anti-α-SMA (1:1000, ET1607-53, HUABIO), anti-GAPDH (1:5000, 60004-1-lg, Proteintech), or anti-α-tubulin (1:1000, AF0001, Byotime) overnight at 4⁏. Binding of the primary antibody was detected by an enhanced chemiluminescence method (BeyoECL Star, P0018A, Byotime) using horseradish peroxidase-conjugated secondary antibodies (goat anti-rabbit IgG, 1:1000, A0208, Beyotime or goat anti-mouse IgG, 1:1000, A0216, Beyotime). The quantification of protein expression was performed using -Image J.

### Statistical analysis

Results were presented as mean ± SD. Differences among multiple groups were analyzed by one-way analysis of variance (ANOVA) and comparison between two groups was performed by unpaired student t-test by using GraphPad Prism version 8.0.0 for Windows (GraphPad Software, San Diego, California USA). A P value of lower than 0.05 was considered statistically significant.

## Author Contributions

MW and CY conceived and coordinated the study. MW wrote the paper. LX conducted the *in vitro* experiments. MW, BT, LX and DH performed the animal experiments. BT measured the concentration of remdesivir and its metabolites. DH, LX and MY performed and analyzed the Western blotting. All authors reviewed the results and approved the final version of the manuscript.

## Acknowledgments

This work was supported by Key Disciplines Group Construction Project of Pudong Health Bureau of Shanghai (PWZxq2017-07), The Three Year Action Plan Project of Shanghai Accelerating Development of Traditional Chinese Medicine (ZY(2018-2020)-CCCX-2003-08) and National Natural Science Foundation of China (81873617) to CY and (81603591) to XL, Scientific Research Foundation of Shanghai Municipal Commission of Health and Family Planning (201740193) to MW.

Disclosures: None.

